# Sleep deprivation triggers somatostatin neurons in prefrontal cortex to initiate nesting and sleep via the preoptic and lateral hypothalamus

**DOI:** 10.1101/2020.07.01.179671

**Authors:** Kyoko Tossell, Xiao Yu, Berta Anuncibay Soto, Mikal Vicente, Giulia Miracca, Panagiotis Giannos, Andawei Miao, Bryan Hsieh, Ying Ma, Raquel Yustos, Alexei L. Vyssotski, Tim Constandinou, Nicholas P. Franks, William Wisden

## Abstract

Animals undertake specific behaviors before sleep. Little is known about whether these innate behaviors, such as nest building, are actually an intrinsic part of the sleep-inducing circuitry. We found, using activity-tagging genetics, that mouse prefrontal cortex (PFC) somatostatin/GABAergic (SOM/GABA) neurons, which become activated during sleep deprivation, induce nest building when opto-activated. These tagged neurons induce sustained global NREM sleep if their activation is prolonged metabotropically. Sleep-deprivation-tagged PFC SOM/GABA neurons have long-range projections to the lateral preoptic (LPO) and lateral hypothalamus (LH). Local activation of tagged PFC SOM/GABA terminals in LPO and the LH induced nesting and NREM sleep respectively. Our findings provide a circuit link for how the PFC responds to sleep deprivation by coordinating sleep preparatory behavior and subsequent sleep.

---

Usually mammals undertake specific behaviors before sleep or as they become drowsy^1–6^. We humans put on bed clothes, lie down and tend to curl up in bed. In other mammals sleep preparatory behaviors, such as building a nest, and going to a nest in a burrow, seem innate. Nesting serves both as a protective environment in which predators can be avoided while sleeping, and provides a thermal microclimate, promoting skin warming that in turn induces NREM sleep^3,7^. In mice, nest building prior to sleep can be initiated by decreasing the activity of dopamine neurons in the midbrain ventral tegmental area^1^. Little else, however, is known about this topic.

Based on lesioning, the prefrontal cortex (PFC) also contributes to general nesting behavior^8^. The PFC links, usually via direct projections from excitatory pyramidal neurons, cognitive operations with other brain regions implementing survival and autonomic processes, such as defensive responses^9^, reward^10^ and selection of behavioral states in response to challenges^11^. There has been intriguing speculation that the neocortex can influence sleep-wake circuitry elsewhere in the brain^12,13^. Because the PFC seems particularly sensitive to sleep deprivation^14–16^, which causes functional connectivity to degrade in the PFC more than in other neocortical areas^15,16^, we hypothesize that this brain region could potentially link sleep pressure, which builds up as wakefulness increases^17^, and sleep preparatory behavior.

Although the delta (0.5 to 4 Hz) oscillations that occur in the NREM sleep EEG probably arise from a neocortex-thalamus dialogue^18^, circuits that drive the actual initiation of sleep are usually regarded as “bottom-up”: hypothalamic, midbrain and brainstem aminergic, glutamatergic and GABAergic neurons control the onset of sleep-wake states^18–27^. In general, behavioral responses to drives such as hunger and thirst are integrated in the hypothalamus, which then influences the neocortex and other brain regions to reinforce the required homeostatic response at a behavioral level^28,29^. Nevertheless, there are examples from the other direction: the PFC can impose behavior via projections to the lateral hypothalamus to limit food intake^30^.

In awake cats, full behavioral NREM sleep can be induced by direct 5-Hz electrical stimulation of the PFC^31–33^, and these observations are quite striking^32^: “When the orbital frontal cortex stimulation was initiated, the animal would stop eating, walk away from the food dish and lie down. If the stimulation was continued for 45 or 50 sec, the animal would go into slow wave sleep and continue to sleep after the stimulation was discontinued. The subject could then be easily aroused by a sensory stimulus”. Further supporting a role for the PFC in inducing NREM sleep, delta waves develop in frontal neocortex^34–36^. In older humans, atrophy of the medial PFC correlates with disrupted NREM slow waves^37^; and frontal cortical areas in rodents and humans have higher amplitude delta wave activity after sleep deprivation, relative to posterior areas^38,39^.

Nearly a third of the neurons in the monkey PFC increase their firing rate during cognitive disengagement (*e.g*. on becoming drowsy), eye closure and sleep^40^. A small percentage of these neurons are likely to be SOM/NOS1 neurons. As assessed by c-FOS expression in mice, nitric oxide synthase (NOS1)-expressing GABA neurons become active throughout the neocortex during sleep deprivation (SD)^41–43^. NOS1 cells form a small subset, 11 to 12%, of somatostatin (SOM, *sst*) neocortical GABA neurons^44–49^. Neocortical and PFC SOM cells induce delta power/slow wave activity and NREM sleep when activated^50^. The circuitry for these effects is not known. There are multiple types of SOM cells^48^. Some inhibit dendrites of pyramidal neurons, whereas the SOM/NOS1/GABA cells have long-range axonal projections within the cortex and their axons can travel through the white matter, but the function of these long-range cells is not known^47,48,51^.

As to be expected, the SOM cells within the PFC are involved in many functions; for example, some PFC SOM cells contribute to reading emotions (affective state discrimination)^52^, others encode fear memories^53^; and locally opto-activating neocortical NOS1 and SOM cells increases cerebral blood flow^54,55^. The *c-fos-* dependent activity-tagging technique allows the SOM/GABA cells that become activated by sleep deprivation to be highlighted from the other functions that these cells carry out. We find using this method that for SOM/GABA neurons in the PFC that become particularly activated after sleep deprivation, their selective reactivation induces nesting and NREM sleep via projections to the lateral preoptic (LPO) and lateral hypothalamus (LH) respectively. This highlights a function for long-range SOM/GABA neurons and provides a direct link between sleep preparatory behavior and global (whole animal) sleep.

## RESULTS

### Local opto-activation of sleep deprivation/recovery sleep-tagged GABA PFC neurons induces nesting and NREM sleep

To functionally investigate GABAergic cells that became active following sleep deprivation (SD) and recovery sleep (RS) in the PFC and visual cortex (VC), we used cell-type selective *c-fos*-based activity-tagging^56,57^, and generated V*gat-PFC-Activity-Tag-ChR2-EYFP* and *Vgat-VC-Activity-Tag-ChR2-EYFP* mice (Fig. 1a; see Online Methods). To avoid confounds with maternal nesting behavior, we used singly housed male mice.

**Figure 1.**
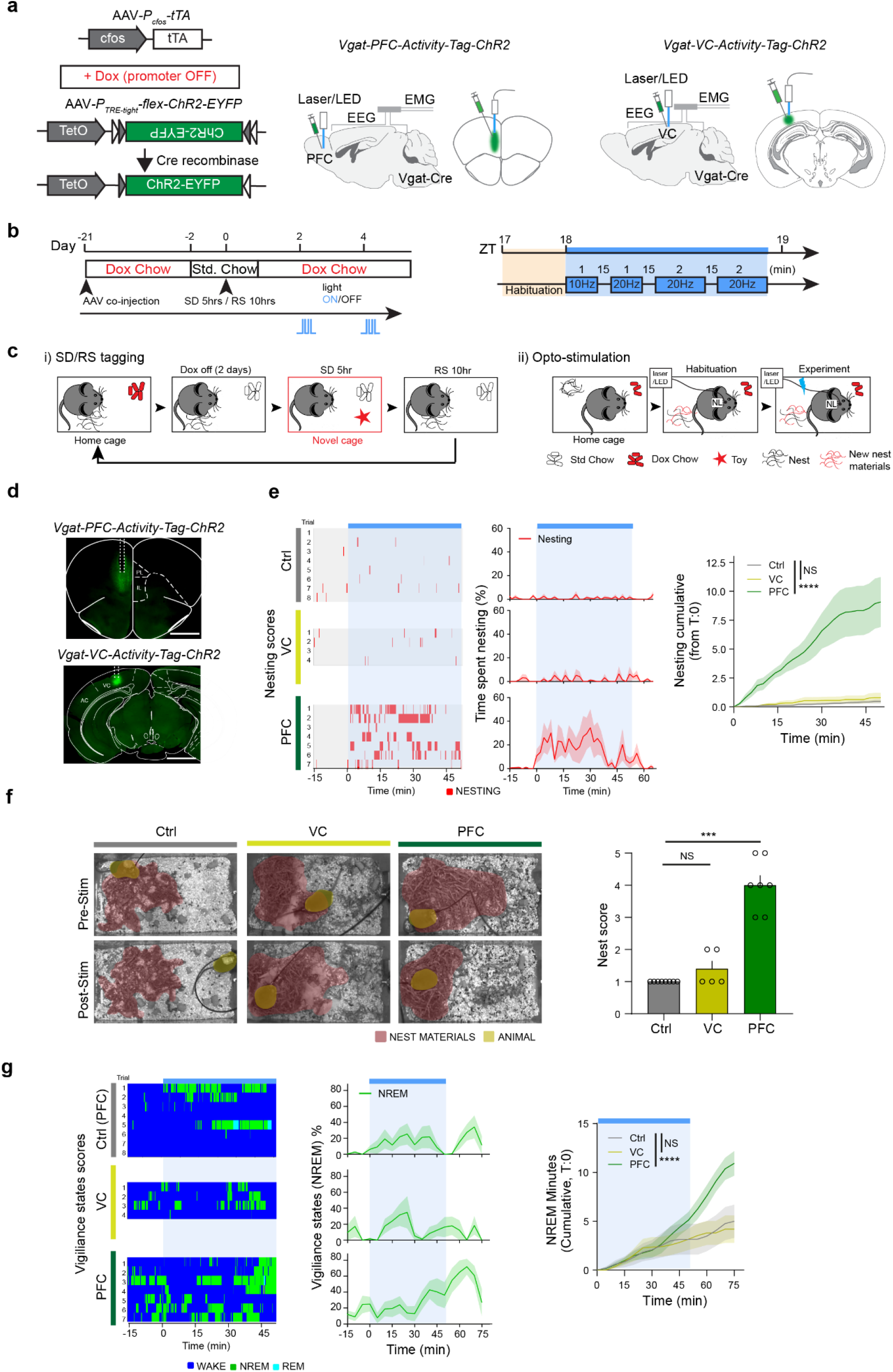
Local activation of activity-tagged PFC-VGAT neurons following sleep deprivation and recovery sleep promotes nesting behavior followed by NREM sleep. **a**, Schematic of (unilateral) activity tagging Vgat-expressing neurons in the PFC and visual cortex (VC) with ChR2-EYFP during sleep deprivation (SD)/recovery sleep (RS). The optic fiber was placed (unilateral) dorsal to the AAV injection site. **b**, Activity-tagging and opto-stimulation protocols. The set of light stimuli was given two days after SD/RS at ZT18. Blue shade indicates the optic-stimulation period. **c**, Schematic illustration of the behavioral experiments as they relate to the activity-tagging and opto-stimulation protocols. Singly housed mice were kept in their home cage with DOX chow before and after SD/RS. SD was performed by moving them to a novel cage with toys. Opto-stimulation was carried out in their home cage, but any existing nest was destroyed, and the material mixed with new nesting materials in order to reduce the habituation period. Food was purposely placed closer to the water bottle and away from the nesting materials to segregate the nest building behavior from food/water seeking behaviors. **d**, Following SD/RS, induced c-Fos-dependent ChR2-EYFP expression was prominently detected in the PFC and VC. Pl, prelimbic; Il, infralimbic; AC, auditory cortex. Scale bar: 1000 μm. **e**, Time course of nesting activity (nesting scores, time spent nesting, cumulative nesting) of opto-stimulated activity-tagged *Vgat-PFC-ActivityTag-ChR2-EYFP* and *Vgat-VC-ActivityTag-ChR2-EYFP* mice. Control (Ctrl) mice included a mixture of PFC-GFP mice and off-Dox non-tagged *Vgat-PFC-Activity-Tag-ChR2-EYFP* mice. “T:0” indicates the starting point of the opto-stimulation block PFC (7 mice), VC (4 mice), Ctrl (4 mice) per group with multiple trials; two-way repeated measures (RM) ANOVA, interaction on PFC and VC group with Ctrl group over time with Bonferroni correction, *F*_25_,_250_ = 0.8662, p = 0.65252 (Ctrl (8 trials) vs VC (5 trials)), *F*_25,325_ = 11.46, p = 1.3089E-31 (Ctrl (8 trials) vs PFC (7 trials)), NS: p>0.05, ****p < 0.0001. Mean (line) ± SEM (shading). Blue shade: Opto-stimulation period. **f**, Pictures of scattered nesting material and nests of *Vgat-PFC-Activity-Tag-ChR2-EYFP* and *Vgat-VC-Activity-Tag-ChR2-EYFP* mice before and after opto-stimulation. Nest material is shaded rose; mouse is shaded olive. PFC (7 mice), VC (4 mice), Ctrl (4 mice) with multiple trials; Unpaired KS test, D = 0.4, p = 0.1282 (Ctrl (8 trials) vs VC (5 trials)), D = 1.0, p = 0.0002 (Ctrl (8 trials) vs PFC (8 trials)), NS: p > 0.05, ***p < 0.001. Mean ± SEM. **g,**Time course of vigilance states (NREM sleep, cumulative NREM sleep) of activity-tagged *Vgat-PFC-Activity-Tag-ChR2-EYFP* and *Vgat-VC-Activity-Tag-ChR2-EYFP* before and during opto-stimulation “T:0” indicates the starting point of the opto-stimulation block. PFC (7 mice), VC (4 mice), Ctrl (4 mice) per group with multiple trials; two-way RM ANOVA, interaction on PFC and VC group with Ctrl group over time with Bonferroni correction, *F*_15_,_150_ = 0.1202, p = 0.99998 (Ctrl (8 trials) vs VC (5 trials)), *F*_15,195_ = 6.023, p = 2.8881E-10 (Ctrl (8 trials) vs PFC (7 trials)), NS: p>0.05, ****p < 0.0001. Mean (line) ± SEM (shading). Blue shade: Opto-stimulation period.

For the activity-tagging experiments, mice were given 5 hours of SD by presenting them with novel objects beginning at Zeitgeber time (ZT) 0 in novel cages. They were then placed back in their home cages and allowed nest building and RS for 10 hours before administering doxycycline (Dox) chow to shut down the activity-tag system (Fig. 1b,ci). SD/RS-tagging produced a strong induction of *ChR2-EYFP* gene expression in GABAergic cells in both the PFC and VC (Fig. 1d & Supplementary Fig. 1a,b). Tagged neurons were analyzed by single-cell PCR in acute slices of PFC prepared from SD/RS-tagged *Vgat-PFC-Activity-Tag-ChR2-EYFP* mice (Supplementary Fig. 1b). None of the SD/RS-tagged cells expressed *vglut1*, which marks pyramidal cells, but all expressed *gad1*, confirming that they were GABAergic. Of these activity tagged GABA cells, about 20% co-expressed the *sst* and *nos1* genes (Supplementary Fig. 1b).

To functionally examine the role of these neocortical GABA cells that became active during SD/RS, a parallel cohort of unilateral *Vgat-PFC-Activity-Tag-ChR2-EYFP* and *Vgat-VC-Activity-Tag-ChR2-EYFP* mice were given SD/RS (Fig. 1a-c). Two days after SD/RS-tagging, unilateral opto-stimulations were given into the PFC or VC and behavior and vigilance states were recorded. We did not know, a priori, the relevant opto-stimulus frequency, so we gave a mixed empirical stimulation protocol of 1-min of 10-Hz light pulses, followed by a 15-min break, then 1 min of 20 Hz followed by a 15-min break and then 2 mins of 20-Hz pulses, amounting to approx. 50 mins of intermittent light stimulation (Fig. 1b). The opto-stimulations were carried out in the home cage, with the mice having been given new nesting materials (Fig. 1cii).

We blindly scored the behavior of the mice in 5-second epochs before and during light stimulation. A notable feature was that during the opto-stimulation, the SD/RS-tagged *Vgat-PFC-Activity-Tag-ChR2*-*EYFP* mice interacted with their nesting material much more so than *Vgat-VC-Activity-Tag-ChR2-EYFP* mice and the groups of control mice (Fig. 1e). In Fig. 1e, the red raster marks indicate when mice were interacting with nesting material *i.e.* pushing and carrying the nesting material; or fluffing the material up, or body wriggling at the center of the nest site, making space for the new nesting material. Most raster scores were in the PFC experimental group during opto-stimulation for individual mice (Fig. 1e); these data are also displayed as “time spent nesting” in the right-hand panel of Fig. 1e. For the SD/RS-tagged *Vgat-PFC-Activity-Tag-ChR2-EYFP* mice, there was a significant accumulation of nesting activity during the 50-min opto-stimulation period. Furthermore, the nests built by the SD/RS-tagged *Vgat-PFC-Activity-Tag-ChR2-EYFP* mice were of higher quality than those built by other groups of mice (Fig. 1f).

During this particular light stimulation protocol both *Vgat-PFC-Activity-Tag-ChR2-EYFP* and *VC-Activity-Tag-ChR2-EYFP* mice stayed awake, and they did not sleep above the baseline of the control mouse groups (Fig. 1g and Supplementary Fig. 1c). However, the opto-stimulated *Vgat-PFC-Activity-Tag-ChR2* mice entered NREM sleep above baseline levels after the opto-stimulation period, coinciding with the time period when nesting behaviour has decreased (Fig. 1g and Supplementary Fig. 1c), whereas the other groups of mice did not. (Examples of individual EEG and EMG spectra for these experiments are shown in Supplementary. Fig. 1c).

We next looked more closely at the opto-stimulation frequencies required to elicit nesting in the SD/RS-tagged *Vgat-PFC-Activity-Tag-ChR2* mice (Supplementary Fig. 2). Following SD/RS-tagging, we gave repetitive 2-minute opto-pulses of either 5, 10 or 20 Hz with 10-min intervals between pulses over a 50-minute period between ZT18 and ZT19 (Supplementary Fig. 2a). All frequencies significantly increased nesting behavior during the opto-stimulation period (Supplementary Fig. 2b). At the end of the stimulation, nesting scores were significantly above baseline for all stimulation frequencies (Supplementary Fig. 2c). For both the 5- and 10-Hz opto-stimulation frequencies, the cumulative amount of NREM sleep gradually increased over the stimulation period, with the biggest cumulative effect seen with 10-Hz stimulation, whereas 20 Hz was ineffective (Supplementary Fig. 2d).

### Metabotropic activation of PFC GABA cells following SD induces NREM sleep

We repeated these SD/RS-tagging experiments using chemogenetics with *Vgat-PFC-Activity-Tag-hM3Dq-mCherry* mice, generated with a small AAV unilateral injection volume into the PFC (Supplementary Fig. 3a,b). Two days following SD/RS tagging, an *i.p.* injection of CNO (1 mg/kg) at ZT18 elicited prolonged nesting behavior and high nest quality scores (Supplementary Fig. 3a & b, c).

We next performed bilateral injections of chemogenetic activity-tag viruses into the PFCs of *Vgat-Cre*, *NOS1-Cre* and *SOM-Cre* mice to generate *Vgat-PFC-Activity-Tag-hM3Dq-mCherry, NOS1*-*PFC-Activity-Tag-hM3Dq-mCherry* and *SOM-Activity-Tag-hM3Dq-mCherry* mice respectively (Fig. 2a and see Online Methods). In all three types of mice, following SD/RS, there was a strong induction of c-fos dependent hM3Dq-mCherry expression in the PFC (Fig. 2a & Supplementary Fig. 4a). CNO (5 mg/kg) was injected *i.p.* 2 days later at ZT18. The total duration of nesting behavior prior to the CNO induced NREM sleep was increased in *Vgat-Cre* and *SOM-Cre* mice, although not in *NOS1-Cre* mice (Supplementary Fig. 3d).

**Figure 2:**
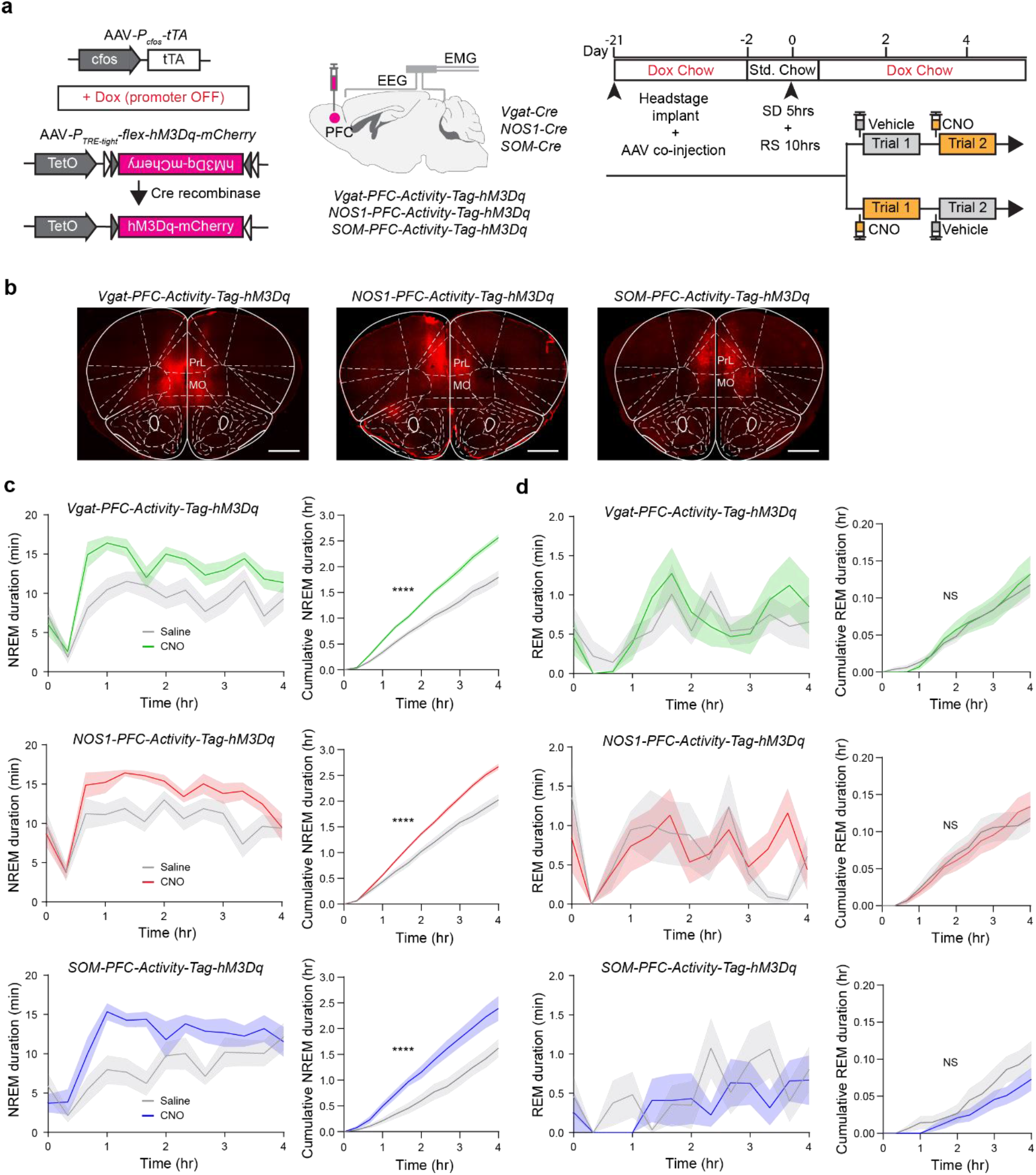
Pharmacogenetic reactivation of activity-tagged GABAergic, NOS1 and SOM neurons in the PFC following sleep deprivation/recovery sleep induces NREM sleep. **a,** Schematic of (bilateral) activity-tagging of Vgat-expressing, NOS1-expressing and SOM-expressing neurons in the PFC with hM3Dq-mCherry during SD/RS; and experimental plan and time course of experiments, showing DOX treatments, SD and RS phases. **b,** Immunohistochemical staining shows that SD/RS induces activity-tagged expression of hM3Dq-mCherry receptor in Vgat, NOS1 and SOM neurons in the PFC. PrL: prelimbic cortex, MO: medial orbital cortex. Scale bar: 1000 μm. **c,** Time course of NREM sleep induction and cumulative NREM amounts evoked by CNO compared with saline injections in *Vgat-PFC-Activity-Tag-hM3Dq*, *NOS1-PFC-Activity-Tag-hM3Dq* mice, and *SOM-PFC-Activity-Tag-hM3Dq* (n=14 (Vgat), n=12 (NOS1), n=10 (SOM) per group; two-way RM ANOVA, Saline vs CNO treatment interaction over time (cumulative NREM sleep) with Bonferroni correction, *F*_12,312_ = 20.26, p = 1.308E-32 (Vgat), *F*_12,252_ = 19.91, p = 2.737E-30 (NOS1), *F*_12,168_ = 7.742, p = 2.318E-11 (SOM). ****p < 0.0001. Mean (line) ± SEM (shading). **d,** Time course of REM sleep induction and cumulative REM amounts evoked by CNO compared with saline injections in *Vgat-PFC-Activity-Tag-hM3Dq* and *SOM-PFC-Activity-Tag-hM3Dq* and *NOS1-PFC-Activity-Tag-hM3Dq* mice (n=14 (Vgat), n=12 (NOS1), n=10 (SOM) per group); two-way RM ANOVA, Saline and CNO treatment interaction over time (cumulative REM sleep) with Bonferroni correction, *F*_12,312_ = 0.2719, p=0.9930 (Vgat), *F*_12,252_ = 0.4959, p=0.9162 (NOS1), *F*_12,216_ = 1.317, p=0.2102 (SOM).NS: p>0.05. Mean (line) ± SEM (shading).

For all three (*Vgat, NOS1, SOM*) groups of mice, sustained NREM sleep was induced above baseline within 1 hour of CNO injection compared with saline (Fig. 2c), REM sleep was unaffected (Fig. 2d). For all three groups of experimental mice, the time course of CNO-evoked NREM sleep was broadly similar (Fig. 2d). [Note: In control mouse strains, CNO (either 1 mg/kg or 5 mg/kg) did not induce NREM sleep or change the amount of wakefulness compared with saline injections (Supplementary Fig. 4b), as we found previously^23,56^]. We also performed SD/RS-tagging on *Vglut2-PFC-Activity-Tag-hM3Dq-mCherry* mice, which after CNO (5 mg/kg) injection evoked no NREM sleep above baseline (Supplementary Fig, 4b). The *vglut2* gene is not expressed in the neocortex, and this serves as a further control for the specificity of the activity-tagging procedure. In summary, subsets of PFC GABA neurons activated by SD/RS can directly induce nesting and NREM sleep and if their activation is sustained metabotropically the tendency to enter NREM sleep persists over several hours.

### Opto-activation of SOM PFC neurons increases nesting and NREM sleep

Moving in to focus on a subset of PFC GABA cells, we SD/RS tagged unilateral *SOM-PFC-Activity-Tag-ChR2-EYFP* mice. Two days after SD/RS-tagging, light stimulation was given into the PFC as 5 blocks of 2 mins of 1-Hz, 5-Hz, or 10-Hz stimulations with 10-mins break between the stimulation blocks (Fig. 3a). The behavioral baseline evoked by light stimulation was obtained on the same animals before the tagging procedure with SD/RS (marked as “pre-tag” in the figure). All frequencies significantly increased nesting behavior during the opto-stimulation period (Fig 3b). At the end of the stimulation, nesting scores were significantly above baseline for all stimulation frequencies (Fig. 3c). For both the 5 and 10 Hz opto-stimulation frequencies, the cumulative amount of NREM sleep gradually increased over the stimulation period, with the biggest cumulative effect seen with 5-Hz stimulation (Fig. 3d). We next investigated the circuitry responsible.

**Figure 3.**
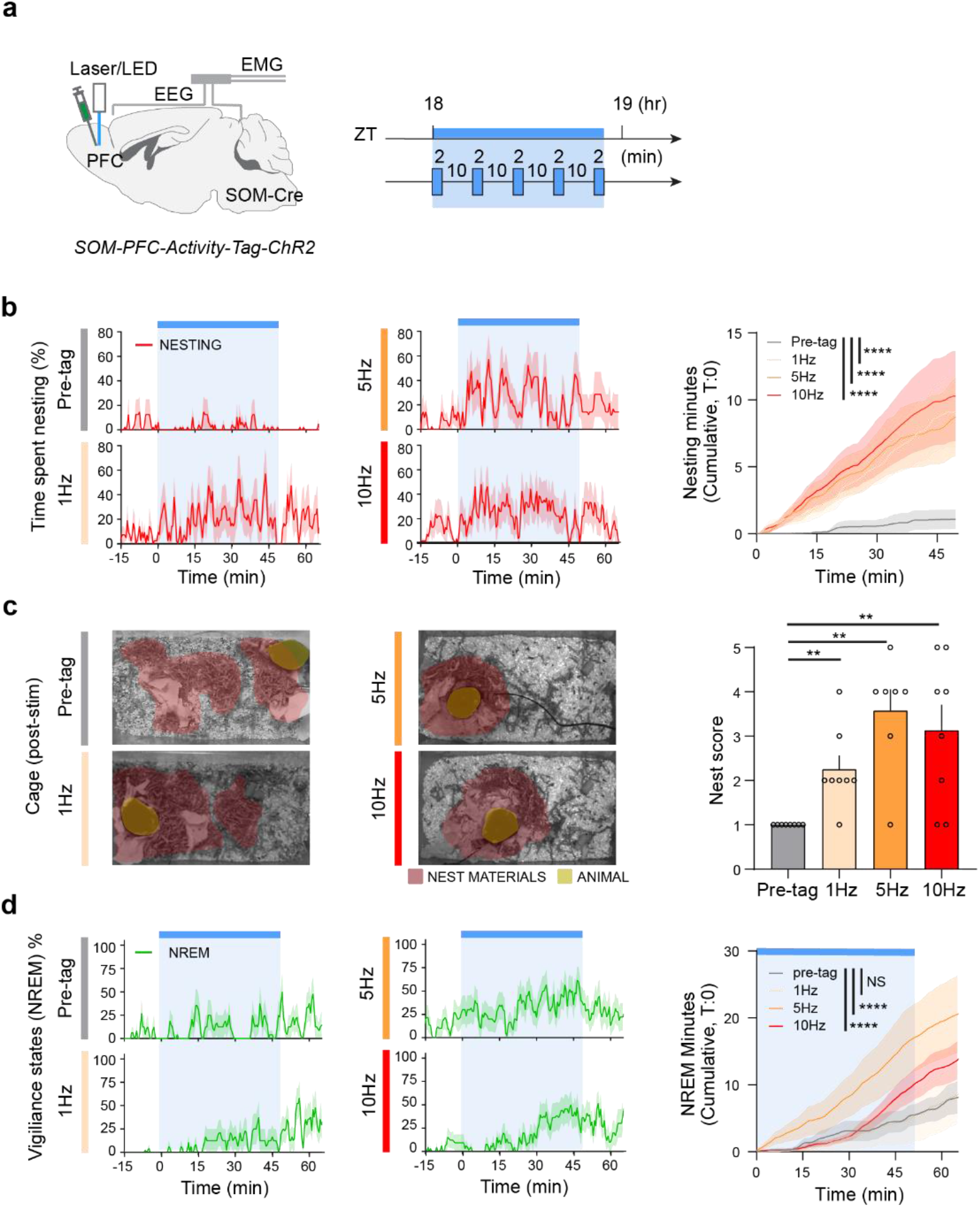
Optogenetic reactivation of activity-tagged SOM neurons in the PFC following sleep deprivation/recovery sleep induces nesting and NREM sleep. **a**, Summary of opto-stimulation protocol for unilateral *SOM-PFC-Activity-Tag-ChR2-EYFP* mice: repetitive blocks of 2 mins stimulation at either 1, 5, and 10 Hz were given with 10 mins intervals over a 50 min block, starting from ZT 18. **b**, Time course of nesting activity (time spent nesting, cumulative nesting) of opto-stimulated SD/RS-tagged *SOM-PFC-Activity-Tag-ChR2-EYFP* mice (n=5 mice each with multiple trials; two-way RM ANOVA, interaction on stimulation group with Pre-tag group over time with Bonferroni correction, *F*_99, 1386_ = 5.692 p = 6.85E-53 (Pre-tag (8 trials) vs 1Hz (8 trials)), *F*_99,1584_ = 4.399, p = 3.49E-37 (Pre-tag (8 trials) vs 5Hz (10 trials)), *F*_99,1683_ = 4.511, p = 5.34E-39 (Pre-tag (8 trials) vs 10Hz (11 trials), ****p < 0.0001. Mean (line) ± SEM (shading). Blue shade: Opto-stimulation period. **c,** Pictures of scattered nesting material and nests of SD/RS-tagged *SOM-PFC-Activity-Tag-ChR2-EYFP* mice after opto-stimulation at either 1, 5, or 10 Hz. Nest material is shaded rose; mouse is shaded olive; and nest scores shown after opto-stimulation. (n=5 mice each with multiple trials; Unpaired KS test, D = 0.875, p = 0.0014 (Pre-tag (8 trials) vs 1Hz (8 trials)), D = 0.875, p = 0.0014 (Pre-tag (8 trials) vs 5Hz (7 trials)),D= 0.75, p = 0.007 (Pre-tag (8 trials) vs 10Hz (8 trials)), **p < 0.01. Mean ± SEM. **d**, Time course of NREM sleep (cumulative sleep) of opto-stimulated SD/RS-tagged *SOM-PFC-Activity-Tag-ChR2-EYFP* mice. (n=5 mice each with multiple trials; two-way RM ANOVA, interaction on stimulation group with Pre-tag group over time with Bonferroni correction, *F*_99, 1386_ = 0.09387, p = 1.0000 (Pre-tag (8 trials) vs 1Hz (8 trials)), *F*_99,1584_ = 3.055, p = 6.3E-20 (Pre-tag (8 trials) vs 5Hz (10 trials)), *F*_99,1683_ = 2.265, p = 9.44E-11 (Pre-tag (8 trials) vs 10Hz (11 trials), NS: p > 0.05, ****p < 0.0001. Mean (line) ± SEM (shading). Blue shade: Opto-stimulation period.

### PFC SOM terminals in LPO induce immediate nesting and a gradual increase of NREM sleep

To map out the circuitry involved in inducing nesting and sleep from PFC GABA/SOM neurons, we SD/RS-tagged unilateral *Vgat-PFC-Activity-Tag-ChR2-EYFP* and *SOM-PFC-ActivityTag-ChR2-EYFP* mice (Fig. 4a – d, Supplementary Fig. 5) and traced out the pulse of ChR2-EYFP protein that was transported into axons during the tagging. In both groups of mice, we found long-range projections of labeled axons from the SD/RS-tagged PFC Vgat and SOM neurons to various areas, including the lateral preoptic (LPO) hypothalamus, the lateral hypothalamus (LH), the medial/lateral habenula (MHb/LHb), the anteriodorsal/medial dorsal (AD/MD) thalamus, the substantia nigra reticulata (SNr) and the periaqueductal grey (PAG) (Fig. 4a, Supplementary Fig. 5). No labeled terminals or fibers were detected in the caudate-putamen (CPu) (Supplementary Fig. 5). In the first instance, we focused on the PO and LH as areas likely to be involved in the sleep and/or nesting behaviors. Both the LPO and LH contained fine ChR2-EYFP positive fibers and punctate structures (Fig 4b).

**Figure 4.**
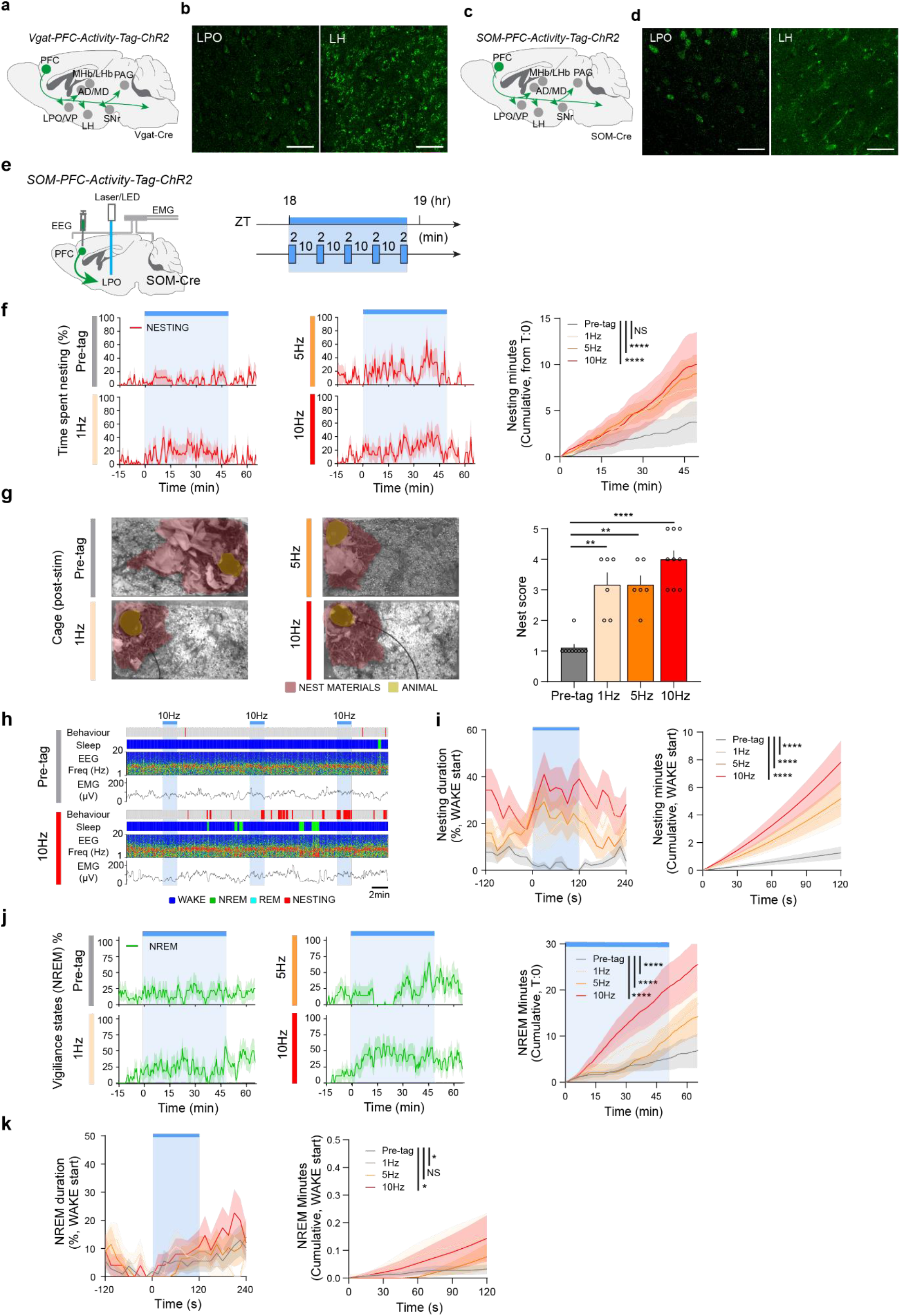
PFC SOM projections to the LPO hypothalamus induce nesting and a gradual increase of NREM sleep. **a,** SD/RS-tagging of PFC Vgat neurons and axons in *Vgat-PFC-Activity-Tag-ChR2-EYFP* mice. Green circle indicates AAV injection area, also illustrated in the left-hand-coronal section. Grey circles and green arrows indicate projection targets **b**, SD/RS-tagged *Vgat-PFC-Activity-Tag-ChR2-EYFP* mice: photographs of *ChR2-EYFP-*positive fibers in the LPO, and LH. Scale bar, 25 μm. **c,** SD/RS-tagging of PFC SOM neurons and axons in *SOM-PFC-Activity-Tag-ChR2-EYFP* mice. Green circle indicates AAV injection area, also illustrated in the left-hand-coronal section. Grey circles and green arrows indicate projection targets. **d**, SD/RS-tagged *SOM-PFC-Activity-Tag-ChR2-EYFP* mice: photographs of *ChR2-EYFP-*positive fibers in the LPO, and LH. Scale bar, 25 μm **e,** Summary of opto-stimulation protocol for the SD/RS-tagged terminals in LPO of *SOM-PFC-Activity-Tag-ChR2-EYFP* mice: repetitive blocks of 2 mins stimulation at either 1, 5, and 10 Hz were given with 10 mins intervals over a 50 min block, starting from ZT 18. **f,** Time course of nesting activity (time spent nesting, cumulative nesting) of opto-stimulated SD/RS-tagged terminals in LPO of *SOM-PFC-Activity-Tag-ChR2-EYFP* mice. (n=4 mice each with multiple trials; two-way RM ANOVA, interaction on stimulation group with Pre-tag group over time with Bonferroni correction, *F*_99, 1386_ = 0.9981, p = 0.48819 (Pre-tag (9 trials) vs 1Hz (8 trials)), *F*_99,1287_ = 2.325, p = 3.95E-11 (Pre-tag (9 trials) vs 5Hz (7 trials)), *F*_99,1485_ = 1.792, p = 6.13E-06 (Pre-tag (9 trials) vs 10Hz (9 trials), NS: p > 0.05, ****p < 0.0001. Mean (line) ± SEM (shading). Blue shade: Opto-stimulation period. **g,** Pictures of scattered nesting material and nests of *SOM-PFC-Activity-Tag-ChR2-EYFP* mice after opto-stimulation of activity-tagged LPO terminals at either 1, 5, or 10 Hz. Nest material is shaded rose; mouse is shaded olive; and nest scores shown after opto-stimulation. (n=4 mice each with multiple trials; Unpaired KS test, D = 0.8889, p = 0.0014 (Pre-tag (9 trials) vs 1Hz (6 trials)), D = 0.8889, p = 0.0014 (Pre-tag (9 trials) vs 5Hz (6 trials)),D= 1.000, p < 0.0001 (Pre-tag (9 trials) vs 10Hz (9 trials), *p < 0.05, **p < 0.01, ****p < 0.0001. Mean ± SEM. **h**, Example traces of EMG and EEG and sleep and behavioral scoring (WAKE, NREM, REM sleep and nesting) from opto-stimulation of activity tagged LPO terminals in *SOM-PFC-Activity-Tag-ChR2-EYFP* mice. Wake is shaded blue, NREM is shaded green, REM is shaded turquois, and nesting behavior is shaded red. Blue shade: Opto-stimulation period (2min). **i,** Nesting behavior duration and accumulation over a 2 min stimulation period of activity tagged LPO terminals in *SOM-PFC-Activity-Tag-ChR2-EYFP* mice which the stimulation started in WAKE period. (n=4 mice each with multiple trials; two-way RM ANOVA, interaction on stimulation group with Pre-tag group over time with Bonferroni correction, *F*_8, 512_ = 7.136, p = 5.68E-09 (Pre-tag (38 trials) vs 1Hz (28 trials)), *F*_8,520_ = 10.02, p = 4.99E-13 (Pre-tag (38 trials) vs 5Hz (29 trials)), *F*_8,488_ = 22.47, p = 2.93E-29 (Pre-tag (38 trials) vs 10Hz (25 trials), ****p < 0.0001. Mean (line) ± SEM (shading). Blue shade: Opto-stimulation period (2min). **j,** Time course of NREM sleep (cumulative sleep) of opto-stimulated SD/RS-tagged terminals in LPO of *SOM-PFC-Activity-Tag-ChR2-EYFP* mice. (n=4 mice each with multiple trials; two-way RM ANOVA, interaction on stimulation group with Pre-tag group over time with Bonferroni correction, *F*_159,2226_ = 1.863, p = 1.78E-09 (Pre-tag (8 trials) vs 1Hz (8 trials)), *F*_159,2067_ = 1.661, p = 1.16E-06 (Pre-tag (8 trials) vs 5Hz (7 trials)), *F*_159,2385_ = 6.401, p = 3.0E-99 (Pre-tag (8 trials) vs 10Hz (9 trials)), ****p < 0.0001. Mean (line) ± SEM (shading). Blue shade: Opto-stimulation period. **k,** NREM sleep duration and accumulation over a 2-min stimulation period of activity tagged LPO terminals in *SOM-PFC-Activity-Tag-ChR2-EYFP* mice which the stimulation started in WAKE period. (n=4 mice each with multiple trials; two-way RM ANOVA, interaction on stimulation group with Pre-tag group over time with Bonferroni correction, *F*_8, 496_ = 2.195, p = 0.02657 (Pre-tag (35 trials) vs 1Hz (28 trials)), *F*_8,480_ = 1.397, p = 0.19518 (Pre-tag (35 trials) vs 5Hz (26 trials)), *F*_8,472_ = 2.22, p = 0.02492 (Pre-tag (35 trials) vs 10Hz (25 trials), *p < 0.05, NS: p > 0.05. Mean (line) ± SEM (shading). Blue-shade: Opto-stimulation period (2min).

The SD/RS-tagged terminals of PFC SOM neurons in the LPO were opto-stimulated at 1, 5 and 10 Hz (Fig. 4e), and the behavior of the mice blindly scored. During the stimulation of terminals over a 50-min period, with 10 minutes intervals between stimuli, the 5- & 10-Hz opto-stimulations evoked an increase of nesting behavior in SD/RS-tagged animals (Fig. 4f). At the end of the stimulation protocol, the nesting scores were ranked significantly higher in the SD/RS-tagged animals compared to the light-stimulated pre-tag baseline group (Fig. 4g). Looking at a finer temporal resolution during an individual 2-min light stimulus of the LPO tagged terminals in which the light stimulus started while the mice were in a WAKE period, the 1-, 5- and 10-Hz opto-stimulations all induced nesting, as assessed by an immediate rate of increase of nesting, with 10 Hz giving the largest effect (Fig. 4h,i).

During opto-stimulation (1, 5 & 10 Hz) of SD/RS-tagged PFC SOM terminals in the LPO area, there was a gradual cumulative increase in NREM sleep above baseline during the 50 minutes of opto-stimulation associated with the 10 Hz stimulations (Fig. 4j). At a 2-min stimulus level, however, although there was a small increase in NREM sleep, there was little correlation with NREM appearance and light stimulus at any frequency (Fig. 4k).

Overall, these results show that selective stimulation of tagged GABA/SOM terminals in the LPO area following sleep deprivation can initiate immediate nesting behavior. Although NREM sleep does increase following light stimulation of LPO terminals, its occurrence is not strongly associated with immediate stimulation of the terminals.

### PFC SOM terminals in the LH induce NREM and REM sleep

As for the LPO experiments described above, following SD, the tagged terminals of PFC SOM neurons in the LH were opto-stimulated at 1, 5 or 10 Hz (Fig. 5a), and the behavior of the mice examined. For all three opto-stimulation frequencies of terminals over a 50-minute period, there was no increase in nesting behavior compared with baseline pre-tagged mice (Fig. 5b); however, when mice did build a nest, the nest quality was scored higher than baseline (Fig. 5c), suggesting that there could be a subtle effect on stimulation of nest building mediated by PFC SOM LH projections.

**Figure 5.**
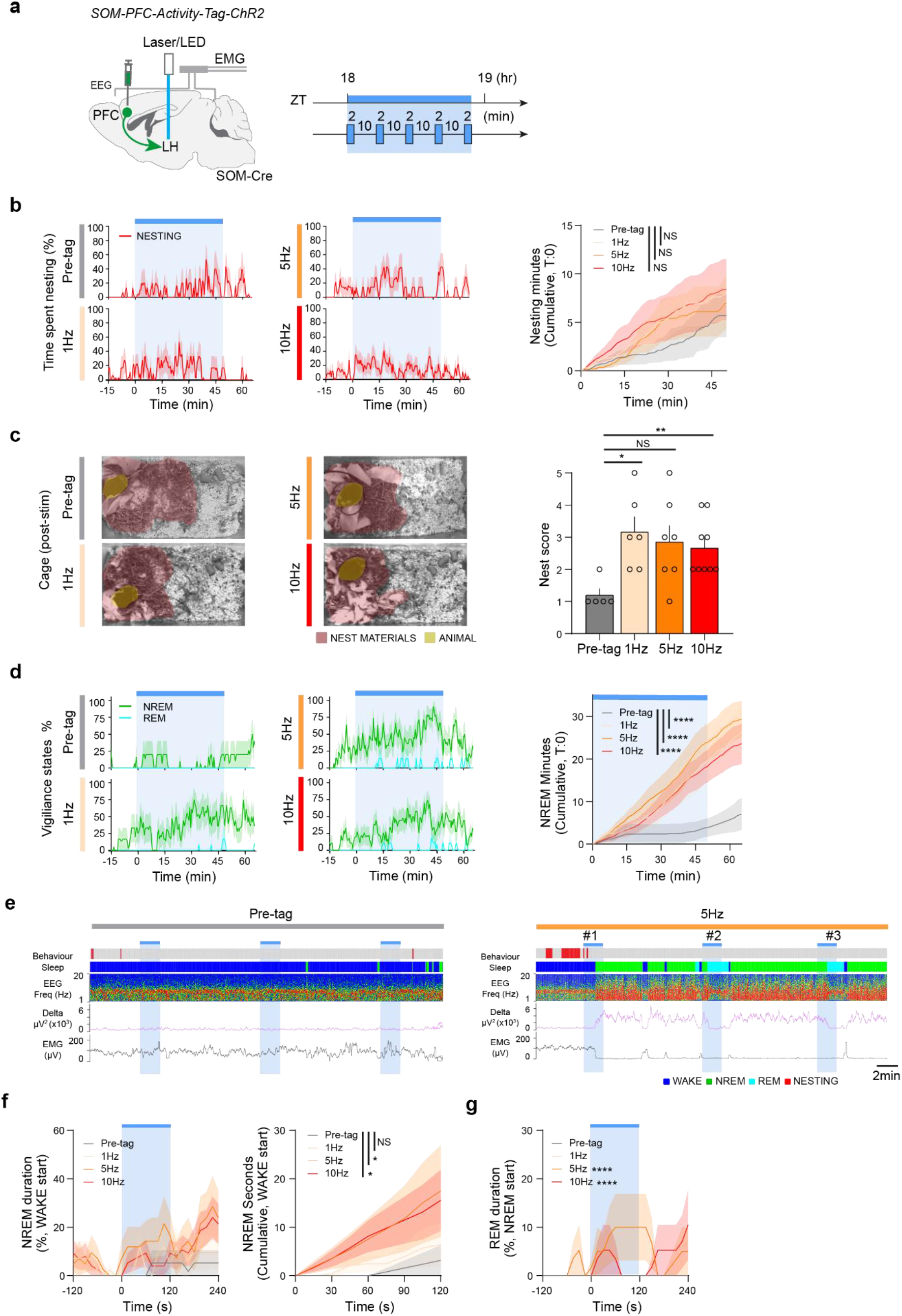
PFC SOM terminals in the LH induce NREM and REM sleep. **a,** Summary of opto-stimulation protocol for the SD/RS-tagged terminals in the LH of *SOM-PFC-Activity-Tag-ChR2-EYFP* mice: repetitive blocks of 2-mins stimulation at either 1, 5, and 10 Hz were given with 10-mins intervals over a 50-min block, starting from ZT 18, which is about 6 hours before lights on. **b**, Time course of nesting activity (time spent nesting, cumulative nesting) of opto-stimulated SD/RS-tagged terminals in LH of *SOM-PFC-Activity-Tag-ChR2-EYFP* mice. (n=4 mice each with multiple trials; two-way RM ANOVA, interaction on stimulation group with Pre-tag group over time with Bonferroni correction, *F*_100, 900_ = 0.9137, p = 0.71154 (Pre-tag (5 trials) vs 1Hz (6 trials)), *F*_100, 1000_ = 0.6973, p = 0.98848 (Pre-tag (5 trials) vs 5Hz (7 trials)), *F*_100, 1200_ = 0.471, p = 0.9999 (Pre-tag (5 trials) vs 10Hz (9 trials), NS: P>0.05. Mean (line) ± SEM (shading). Blue shade: Opto-stimulation period. **c,** Pictures of scattered nesting material and nests of *SOM-PFC-Activity-Tag-ChR2-EYFP* mice after opto-stimulation of SD/RS-tagged LPO terminals at either 1, 5, or 10 Hz. Nest material is shaded rose; mouse is shaded olive; and nest scores shown after opto-stimulation. (n=4 mice each with multiple trials; Unpaired KS test, D = 0.8, p = 0.0303 (Pre-tag (5 trials) vs 1Hz (6 trials)), D = 0.6571, p = 0.0783 (Pre-tag (5 trials) vs 5Hz (7 trials)),D= 0.8, p = 0.01 (Pre-tag (5 trials) vs 10 Hz (9 trials), NS> 0.05, *p < 0.05, **p < 0.01. Mean ± SEM. **d,** Time course of NREM and REM sleep (cumulative sleep) of opto-stimulated SD/RS-tagged terminals in LH of *SOM-PFC-Activity-Tag-ChR2-EYFP* mice. (n=4 mice each with multiple trials; two-way RM ANOVA, interaction on stimulation group with Pre-tag group over time with Bonferroni correction, *F*_160, 1440_ = 6.014, p = 1.92E-81 (Pre-tag (5 trials) vs 1Hz (6 trials)), *F*_160, 1600_ = 10.96, p = 5.5E-166 (Pre-tag (5 trials) vs 5Hz (7 trials)), *F*_160, 1920_ = 4.401, p = 5.67E-56 (Pre-tag (5 trials) vs 10Hz (9 trials), NS: P>0.05, ****p < 0.0001. Mean (line) ± SEM (shading). Blue shade: Opto-stimulation period. **e**, Example traces of EMG and EEG and sleep and behavioral scoring (WAKE, nesting, NREM and REM sleep) from opto-stimulation of SD/RS-tagged LH terminals in *SOM-PFC-Activity-Tag-ChR2-EYFP* mice. Wake is shaded blue, NREM is shaded green, REM is shaded turquois, and nesting behavior is shaded red. Blue shade: Opto-stimulation period (2min). **f,** NREM sleep duration and accumulation over a 2-min stimulation period of SD/RS-tagged LH terminals in *SOM-PFC-Activity-Tag-ChR2-EYFP* mice, which the stimulation started in WAKE period (WAKE -> NREM). (n=4 mice each with multiple trials; two-way RM ANOVA, interaction on stimulation group with Pre-tag group over time with Bonferroni correction, *F*_8, 248_ = 0.4349, p = 0.89945 (Pre-tag (19 trials) vs 1Hz (14 trials)), *F*_8, 248_ = 2.252, p = 0.02443 (Pre-tag (19 trials) vs 5Hz (13 trials)), *F*_8, 352_ = 2.154, p = 0.03047 (Pre-tag (19 trials) vs 10Hz (24 trials), NS: P>0.05, *p < 0.05. Mean (line) ± SEM (shading). Blue shade: Opto-stimulation period. **g,** REM sleep duration and accumulation over a 2 min stimulation period of SD/RS-tagged LH terminals in *SOM-PFC-Activity-Tag-ChR2-EYFP* mice which the stimulation started in NREM period (NREM -> REM). (n=4 mice each with multiple trials; Two-tailed non-parametric t-test with Bootstrap REM sleep duration (mean) during Opto-stimulation against before Opto-stimulation, U=20201, p > 0.9999 (Pre-tag (NREM -> REM: 7 trials)), U = 20201, p > 0.9999 (1Hz (NREM -> REM: 15 trials)), U = 8472, p < 0.0001 (5Hz (NREM -> REM: 20 trials), U = 11457, p< 0.0001 (10Hz (NREM -> REM: 19 trials). ****p < 0.0001. Mean (line) ± SEM (shading). Blue shade: Opto-stimulation period (2 min).

On the other hand, the 50-min opto-stimulation period of SD/RS-tagged LH terminals at 1, 5 and 10 Hz caused a strong increase in cumulative NREM sleep (Fig. 5d). Looking at a finer temporal resolution, over a 2-min stimulation period, and using the 5-Hz stimulation experiments as an example, there was an immediate increase in the NREM sleep induction from wakefulness (Fig. 5e, two examples are sections of the trace marked #1 and #2 in the right-hand block of traces). Quantitatively, over a particular period of 2-min light stimulation of SD/RS-tagged LH terminals during wakefulness, there was an increase in NREM sleep (Fig. 5f). Additionally, if the mice were already in NREM sleep, further opto-stimulation of SD/RS-tagged LH terminals induced transient REM sleep (Fig. 5e, examples #2 & #3 in the traces; and Fig. 5g). Thus, following SD, PFC SOM neurons can directly trigger sleep via projections to the LH.

## DISCUSSION

After sleep deprivation, mice either go to a prebuilt nest, or if there is no nest, they build one before starting their recovery sleep. We have found that such sleep preparatory behavior and subsequent sleep can be induced top-down from the PFC by GABAergic projections to the hypothalamus. This is consistent with anterograde and retrograde tracing studies that demonstrated afferents between the PFC and LPO and LH areas^58–60^, and provides one function for the mysterious long-range SOM/NOS1/GABA neurons^47,51^. Unilateral opto-stimulation of the sleep-deprivation tagged PFC SOM/NOS1/GABA cells intermittently at 5 to 10 Hz was enough to induce nesting behavior and the accumulation of NREM sleep while the nesting was ongoing. Similarly, sustained metabotropic stimulation of SOM/NOS1/GABA cells in the PFC initiated nesting and was then sufficient to maintain global NREM sleep. Selective opto-stimulation of the sleep deprivation-tagged terminals originating from PFC SOM cells in the LPO and LH induced nesting and NREM sleep respectively. The dual nesting and sleep-induction mechanism, SOM-PFC to LPO, SOM-PFC to LH, could be a way of ensuring sleep takes place in the nest. We envisage that there are parallel routes to sleep induction which reinforce one another. Nesting provides a thermal microclimate, promoting skin warming, which in turn promotes NREM sleep induction^3,7^ via circuitry from receptors in the skin, to neurons in the MPO hypothalamus^56^.

The building of nests by mammals and birds seems mostly an innate behavior, and is a stored series of intricate and sequential motor skills^61^. Little is known about the circuitry regulating nesting in rodents. From lesioning studies, nesting partially requires the PFC^8^. Nesting behavior prior to sleep can be initiated by inhibiting dopamine neurons in the ventral tegmental area (VTA)^1^. For maternal nest building, the instinct partly depends on circuitry in the medial preoptic (MPO) hypothalamus and GABA projections from the arcuate hypothalamus to the MPO^62^; chemogenetic inhibition of GABAergic MPO neurons reduces, and ablation of MPO galanin neurons abolishes, maternal nest-building^63,64^. The LPO area, which is similar to MPO, is highly heterogeneous, and contains multiple cell types and circuits (regulating, for example, sleep, body temperature, blood osmolarity and heart rate^65^), could output to similar circuitry governing nest building.

Following sleep deprivation, the PFC to LH projection could induce NREM sleep by inhibiting arousal-promoting GABA and orexin neurons in the LH^66,67^; this would be a similar mechanism to how VTA GABA neurons induce NREM sleep by sending their GABAergic projections to the LH^23,68,69^. Sustained local stimulation of this projection could disinhibit REM sleep-promoting MCH/GABA cells^70^. This could explain the phenomenon we found that once the mice are already in NREM sleep, further opto-stimulation of the tagged LH terminals induces REM sleep (Figure 5e,g).

There are likely to be several sleep-related mechanisms at play in the neocortex. One is the sleep preparatory behavior and sleep mediated by the PFC to hypothalamus projections reported here. But local sleep could also be induced without nesting behavior by NOS1/SOM neurons in other areas of the neocortex. Excitability throughout the neocortex increases with time spent awake^71^. In awake and behaving animals, local delta NREM-like oscillations develop in different regions of the neocortex following use-dependent activity^72–76^. It is well established that neocortical NOS1-expressing cells become active throughout the neocortex during sleep deprivation^41,42^. Other researchers found that chemo- and optogenetic activation of SOM neurons in the mouse frontal and motor cortex enhanced delta power/slow wave activity in the EEG^50^, and chemogenetic activation of SOM neurons induced NREM sleep^50^, but the circuit mechanisms were not investigated^50^ – the sleep was hypothesized to arise from local dendritic inhibition of pyramidal neurons^50^. Indeed, in slow wave sleep, neocortical SOM cells are active just before the down states of pyramidal cells^77^. When NOS1 was removed selectively from SOM cells, the lack of nitric oxide did not affect occurrence of NREM sleep but reduced its delta power^78^ *i.e.* nitric oxide produced in SOM/GABA cells is needed for the depth of sleep (degree of neocortical synchronization at delta frequencies). This could be regulated by layer V pyramidal neurons. Genetically silencing these neurons blocks the increase in EEG delta power following SD without changing the amount of sleep^79^. By contrast, the SOM/GABA component could be needed for sleep itself.

A picture of sleep-wake regulation is emerging in which there is no single master network generating sleep and wake. Circuitry that senses the need to sleep could be distributed throughout the brain. After sleep deprivation, many brain regions have enhanced activity, as seen by immediate early gene expression^80–82^. The LPO hypothalamic area, for example, has GABAergic neurons that become excited during sleep deprivation/recovery sleep^83^. *cfos*-dependent activity-tagging shows that these LPO neurons are sufficient to sense the sleep drive and promote NREM sleep^57^; reactivation of LPO galanin neurons following sleep deprivation induces NREM sleep^57,84^. Other areas include the VTA. Lesioning VTA GABA neurons abolishes the ability of mice to catch up on lost sleep after sleep deprivation^85,86^. Each area could respond to its own need for the replenishment provided by sleep.

In conclusion, it is important to place sleep in an ethological perspective^2,4^. Our work shows that following sleep deprivation, behavioral preparation for sleep is linked to some of the sleep-induction circuitry. In the case of the PFC, with its role in executive function and planning, a direct connection to hypothalamic centers to initiate sleep preparation, and to help reinforce global sleep, could be a survival advantage to ensure the animal is in a safe place prior to sleeping.

## Methods

### Mice

All experiments were performed in accordance with the UK Home Office Animal Procedures Act (1986), and approved by the Imperial College Ethical Review Committee. The following types of mice were used: *Vgat-ires-Cre: Slc32a1^tm2(cre)Lowl^/J* mice (JAX labs Stock 016962), kindly provided by B.B. Lowell^87^; *Nos1-ires-Cre^tm1(cre)Mgmj^/J* (JAX labs stock 017526) kindly provided by M. G. Myers^88^, SOM-ires-Cre: Sst^tm2.1(cre)Zjh^/J (JAX labs stock 013044) kindly provided by Z. J. Huang^89^, VGlut2-ires-Cre: Slc17^a6tm2(cre)Lowl^/J (JAX labs stock 016963) kindly provided by B. B. Lowell^87^ and *C57Bl/6J* mice (supplied by Charles River UK). All mice used in the experiments were male and congenic on the C57Bl/6J background. Mice were maintained on a 12 hr:12 hr light:dark cycle at constant temperature and humidity with *ad libitum* food and water.

### AAV transgene plasmids & AAV preparation

We have described several of the plasmids previously: pAAV-cFos-tTA-pA (Addgene plasmid #66794)^57^, pAAV-P_TRE-tight_-flex-hM3Dq-mCherry (Addgene plasmid #115161)^56^; and pAAV-flex-EGFP^90^. To make *pAAV-ITR-P_TRE-tight_-flex-ChR2-EYFP-WPRE-pA-ITR*, the backbone *pAAV-ITR-_PTRE-tight_-flex* (4.7kb) was purified from *pAAV-ITR-P_TRE-tight_-flex-hM3Dq-mCherry-WPRE-pA-ITR*, after double digestion with AscI + NheI and gel purification. The insert, *ChR2-EYFP*, was taken from Addgene plasmid number 20298 (humanizedChR2 with H134R mutation fused to EYFP, gift of Karl Deisseroth), after double digestion with AscI + NheI and gel purification. Both fragments were ligated to produce the final construct *pAAV-ITR-PTRE-tight-flex-ChR2-EYFP-WPRE-pA-ITR* (to be deposited at Addgene).

To produce AAV, the adenovirus helper plasmid *pFΔ6*, and the AAV helper plasmids *pH21* (AAV1), *pRVI* (AAV2), and the *pAAV* transgene plasmids were co-transfected into HEK293 cells and the subsequent AAV particles harvested on heparin columns^91^.

### Stereotaxic surgery

One week before surgery, mice were placed on 200 mg/kg doxycycline (Dox) (Envigo TD.09265) chow. Stereotaxic virus injections were performed using an Angle Two apparatus (Leica) linked to a digital brain atlas (Leica Biosystems Richmond, Inc.) and a stainless steel 33-gauge/15-mm/PST3 internal cannula (Hamilton) attached to a glass syringe (10 μl-Hamilton #701). Two types of AAV, e.g. *AAV-P_cFos_-tTA* and *AAV-P_TRE-tight_-flex-hM3Dq-mCherry* or *AAV-P_cFos_-tTA* and *AAV-P_TRE-tight_-flex-ChR2-EYFP* were freshly mixed in a 1:1 ratio just before the surgery. Unless otherwise specified, virus was bilaterally injected at 0.1 μL min^−1^, with up to two injections totalling 0.5 μl for the PFC and 1 μl for the VC. For unilateral injections for the chemogenetic AAV experiments, a single injection of 0.25 μl was made to the left hemisphere of the PFC. The injection co-ordinates were PFC: ML (±0.4 mm), AP (2.1 mm), DV (−2.45 mm); and VC ML (±2.38 mm), AP (−2.54 mm), DV (−0.92 mm).

For the optogenetics experiments, we first injected the AAV mixture unilaterally (for PFC/VC) and bilaterally (for LPO/LH), and then implanted a mono-fiber optic cannula (I.D. 200 μm, 0.37 NA, Thorlabs Cat# FT200EMT) unilaterally directly above the following co-ordinates: LH: ML (±1.0 mm), AP (−1.4 mm) DV (5.16 mm) and LPO: ML (± 0.75 mm), AP (4.0 mm), DV (5.15 mm).

For sleep recordings, two EMG wire electrodes were inserted in the neck extensor muscles and two EEG screw electrodes were placed at ML (−1.5 mm), AP (+1.5 mm) and ML (−1.5 mm), AP (−2.0 mm) relative to Bregma. A third EEG electrode was placed at ML (+1.5 mm), AP (−2.0 mm) for optogenetic recording. All instrumented mice were single-housed to avoid damage to the head-stage, and were allowed to recover and for the viral transgenes to adequately express for at least 3 weeks.

### Activity-Tag behavioral protocols and controls

Two AAVs, *AAV-P_cFos_-tTA* and *AAV*-*P_TRE-tight_-flex-hM3Dq-mCherry*, were bilaterally co-injected into the PFC of *Vgat-Cre* or *Nos1-Cre* or *Som-Cre* mice to generate *Vgat-PFC-Activity-Tag-hM3Dq-mCherry*, *Nos1-PFC-Activity-Tag-hM3Dq-mCherry* and *SOM-PFC-Activity-Tag-hM3Dq-mCherry* mice respectively (Fig. 1a). To repress the activity-tagging system, mice were maintained on Dox chow for one week prior to the surgery and at least 3 weeks post-surgery (Fig. 1b). Prior to sleep deprivation (SD), mice were taken off Dox for 2 days, and then given 5 hours of SD by introducing novel objects beginning in the novel cage at the start of the “lights on” (ZT 0) period. Mice were gently placed back into their home cages, and allowed recovery sleep (RS) for 10 hours (Fig. 1b). After this, the mice were placed back on Dox chow (Fig. 1b). Mice were habituated to the Neurologger 2A EEG recording devices for at least 2 days before the SD/RS was performed. During this time, 24 hr EEG/EMG baseline recording was obtained, and SD/RS was monitored and confirmed offline. Any mice which failed to show 5hr of clear SD and an RS accompanied with a delta power increase were discounted from the chemogenetic or optogenetic experiments.

### EEG/EMG recordings and analysis

EEG and EMG signals were recorded using Neurologger 2A devices^92^ at the sampling rate of 200 Hz and the data was visualized with Spike 2 software (Cambridge Electronic Design, Cambridge, UK). The EEG signals were high-pass filtered offline at 0.5Hz (−3dB) and EMG signals were band-pass filtered offline at 5-45 Hz (−3dB). To define the vigilance states of WAKE, NREM, REM sleep, delta power (0.5-4.5 Hz), theta power (5-10 Hz) and theta/delta (T:D) ratios were calculated. Automated sleep scoring was performed using a Spike 2 script and the results were manually corrected.

### Chemogenetics

Two days after RS, clozapine-N-oxide (CNO) (4936, Tocris, dissolved in saline, 1 mg/kg and 5 mg/kg) or saline was injected intraperitoneally (i.p.) at ZT 18 (i.e. mid-“lights off” period) and the vigilance states recorded (Fig. 2a). Mice were split into random groups that received either saline (day 1) or CNO (day 2) or CNO (day 1) or saline (day 2) injections at the same circadian time. Mice were habituated to the Neurologger 2A devices at least 1 hour before ZT18 (*i.p.* injection T:0).

### Optogenetics

Mice were allowed to habituate to the Neurologger 2A devices minimally 2 days before the SD/RS was performed. Optogenetic stimulations were generated by 473 nm diode-pumped solid-state (DPSS) laser with fiber coupler (Shanghai Laser, BL473T3T8U-100FC, Shanghai Laser & Optics Century, China) or 465 nm Doric Connectorized LED (CLED_465, Doric Lenses, Canada). Stimulation protocols were programmed and controlled using Signal software (Cambridge Electronic Design, UK) and Micro1401 (CED, UK) for laser, and Doric Neuroscience Studio v5.3.3.14 (Doric Lenses, Canada). Laser/LED power was kept in the range of 2-5 mW at the tip of the optic fiber.

Opto-stimulation was carried out during ZT 18 (the mid-lights-off period in the animal house) *i.e*. at the time when mice were most active and least likely to build a nest or sleep. Before starting the stimulation protocol, all mice were habituated for at least 30 minutes to the environment. For control to SD/RS tagged *Vgat-PFC-Activity-Tag-ChR2* and *Vgat-VC-Activity-Tag-ChR2* mice, we used *Vgat-PFC-Activity-Tag-ChR2* mice that had had DOX removed from their diet for the same time duration as the paired experimental cohorts but that had received no SD/RS, and *Vgat-PFC-GFP* mice (AAV-flex-EGFP were injected into PFC of *Vgat-Cre* mice). Both groups were implanted with optic fibers and given light stimulation as the same manner to experimental groups. Results from both groups of controls were pooled.

### Behavioral analysis and nest scoring

All behavior was monitored by video camera which was placed above the test cage, and analyzed offline after the experiments. All evaluation was carried out on pre-blinded recording data by more than one experimenter. The difference was carefully reviewed and corrected before unblinding. Videos were synchronized with stimulation protocols. Video nesting behavior over time was scored using Behavioral Observation Research Interactive Software (BORIS)^93^ and aligned with sleep scoring in Spike 2. Nesting behavior was defined as pushing and carrying the nesting material; or fluffing the material up, or body wriggling at the center of the nest site, and making space for the new nesting material.

Before the initial habituation period (start at ZT17) of optogenetic experiment, all previously existed nest was removed from home cage of the test mice and placed 8g of mixture of old and new shredding papers, away from food and water. Baseline nest condition was remotely checked 5-10 mins before ZT18 (opto-stimulation T:0) without disturbing the test mice. For the chemogenetics (unilateral, low-dose) experiments, the cage was prepared 30 min to 1 hour before i.p. injection at ZT18, and monitored by the overhead video camera for 5 hours.

We evaluated nest scores 1 hr after opto-stimulation period offline (at ZT19) by adapting a five-point scale^1^.

1. Nest materials not noticeably touched (<10% change from baseline)
2. Nest materials partially gathered (10-50% change from baseline)
3. Nest materials sorted and gathered, but some are spread around the cage (50-90% change from baseline)
4. Nest materials sorted and gathered, identifiable but flat.
5. A perfect or near perfect nest with a crater.

### Immunohistochemistry and imaging

Mice were transcardially perfused with 4% paraformaldehyde (Thermo scientific) in phosphate buffered saline (PBS). Brains were removed and 35-μm-thick coronal sections were cut using a Leica VT1000S vibratome. Free-floating sections were rinsed once in PBS and processed for epitope retrieval by incubating sections in 0.05% Tween20 in 10 mM sodium citrate buffer (pH6.0) at 80-85 °C for 30min. Sections were allowed to cool down to the room temperature (RT), then washed 3 times with PBS for 10 min. Sections were blocked with 20% goat serum (NGS: Vector)/0.2% Triton X-100/PBS solution for 1 hr at RT, and incubated with primary antibody at adequate dilution in 2% NGS/0.2% Triton X /PBS solution for overnight at 4 °C. Incubated slices were washed three times in PBS for 10 min at RT, and incubated with a secondary antibody (Molecular Probes) at adequate dilution in 2% NGS/ 0.2% Triton X/PBS solution for 1.5 hr at RT. Slices were washed 3 times with PBS for 10 min at RT, and incubated in Hoechst 33342 (Life Technologies) at 1:5000 in PBS for up to 10 mins at RT. After a double wash in PBS, slices were mounted on slides with ProLong Gold Antifade Reagent (Invitrogen). Primary antibodies: rabbit anti-GFP (Invitrogen, A6455, 1:1000), chicken anti-GFP (Abcam, ab13970, 1:1000), mouse-anti-mCherry (Clontech, 632543 1:1000). Secondary antibodies: Alexa Fluor-488 goat anti-chicken (Invitrogen, A11039, 1:500), Alexa Fluor-488 goat anti-rabbit (Invitrogen, A11008, 1:500), Alexa Fluor-594 goat anti-mouse (Invitrogen, A11005, 1:500). Image were taken with Axiovert 200M Inverted widefield microscope (Zeiss) and Leica SP8 Inverted confocal microscope.

### Single-cell RT-PCR of activity-tagged neurons

*Vgat-PFC-Activity-Tag-ChR2-EYFP* mice that had undergone the SD/RS procedure and that had been placed back onto Dox were euthanized by cervical dislocation. The brain was quickly removed and placed into cold oxygenated *N*-Methyl-*D*-glucamine (NMDG) solution (in mM): NMDG 93, HCl 93, KCl 2.5, NaH_2_PO_4_ 1.2, NaHCO_3_ 30, HEPES 20, glucose 25, sodium ascorbate 5, Thiourea 2, sodium pyruvate 3, MgSO_4_ 10, CaCl_2_ 0.5. Coronal brain slices (170-μm thickness) encompassing the PFC were obtained using a vibratome (Vibrating Microtome 7000smz-2; Campden Instruments LTD, UK). Slices were transferred to a submersion chamber and continuously perfused at a rate of 1-2 ml/ min with fully oxygenated aCSF at room temperature. Activity-tagged neurons were identified by their EYFP signal under fluorescence illumination (LED4D, Thorlabs, coupled to YFP excitation filter). Neuronal contents were aspirated into borosilicate glass capillaries and expelled into cell lysis/DNase I solution. The cDNA synthesis was performed using the Single-Cell to-CT Kit (Ambion) and qPCR was performed using the TaqMan Gene Expression Assay system (Applied Biosystems machine, Foster City, USA). The mouse TaqMan assay probes were designed by, and purchased from, Invitrogen: m18srRNA, Mm03928990_g1; mNpy Mm01410146_m1, mNos1: Mm01208059_m1; mSst: Mm00436671_m1. mVglut1: Mm00812886_m1, mGad1: Mm00725661_s1.

### Quantification and Statistical Analysis

Prism v8.4.2 was used for statistical analysis. Data collection and processing were randomized or performed in a counter-balanced manner. P values are shown when they are less than 0.05 (*p < 0.05, **p < 0.01, ***p < 0.001, ****p < 0.0001). For nesting behavior and NREM/REM sleep analysis, 2 way-repeated measure ANOVA with Bonferroni correction was used. For nest score, unpaired-KS test was used. Mice were excluded from the analysis if the histology did not confirm AAV transgene expression in the PFC or VC. While experimenters were not blinded to treatments, data analysis were carried out blindly.

## AUTHOR CONTRIBUTIONS

K.T., X.Y. N.P.F., and W.W. conceived and designed the experiments; K.T., X.Y., A.M., B.A.C., P.G., M.V., B.H., Y.M., G.M., A.M., W.S.T. and R.Y. performed the experiments and data analysis; A.L.V. provided the Neurologger 2A devices; T.C. contributed electrical engineering advice and co-supervised B.H, N.P.F. and W.W. contributed to the data analysis, and co-supervised the project; K.T, X.Y., N.P.F., and W.W. co-wrote the paper.

## ACKNOWLEDGEMENTS

This work was supported by the Wellcome Trust (107839/Z/15/Z, N.P.F. and 107841/Z/15/Z, W.W), the UK Dementia Research Institute at Imperial College (W.W and N.P.F.), an MRC UK DRI studentship (A.M.), an Imperial College Schrodinger Scholarship (G.M.), an Imperial College/China Scholarship scheme (Y.M.); and EPSRC studentship, EPSRC Centre for Doctoral Training in Neurotechnology for Life and Health (B.H.). The facility for Imaging by Light Microscopy (FILM) at Imperial College London is part-supported by funding from the Wellcome Trust (104931/Z/14/Z) and BBSRC (BB/L105129/1).

## Supp. Figures.

**Supplementary. Fig. 1.**
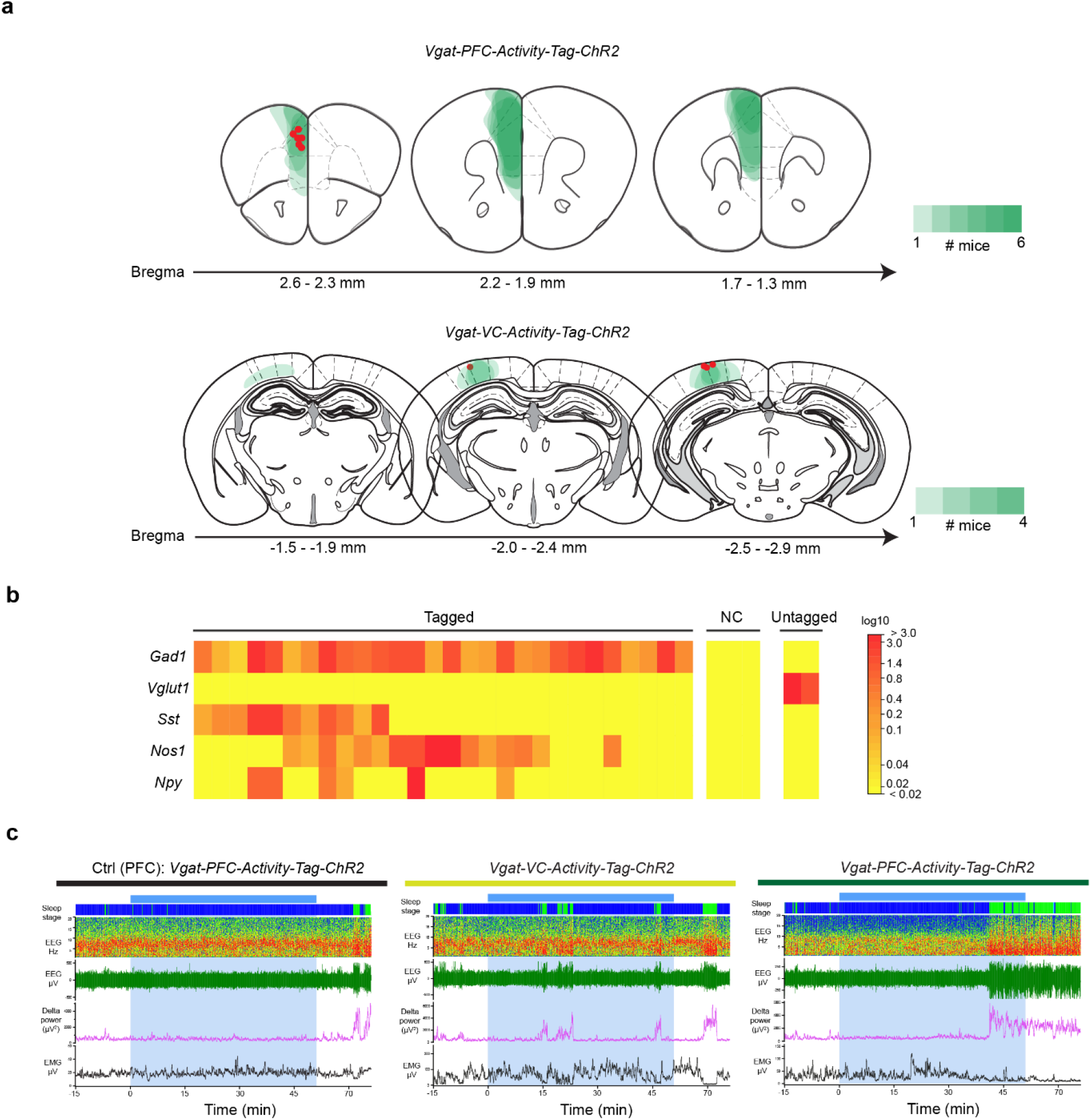
(supports Fig. 1). **a,** Map of ChR2-EYFP gene expression induced following SD/RS in *Vgat-PFC-Activity-Tag-ChR2-EYFP* and *Vgat-VC-Activity-Tag-ChR2-EYFP* mice. Photomicrograph inset shows tagged neurons in the PFC. The schematic diagrams summarize the extent of gene expression over 6 animals in the PFC and 4 mice in the VC (the intensity of the green indicates the extent of overlap (overlay) of gene expression). Red dot marks the position of the optic fiber tract i.e. where the cortex is being stimulated **(Supports Figure 1)**. **b**, Single-cell PCR from visually identified activity-tagged neurons in acute PFC slices prepared from SD/RS-tagged *Vgat-PFC-Activity-Tag-ChR2-EYFP* mice. Heat map, red is strongest expression; yellow is weaker. NC, negative control. “untagged cells” were pyramidal neurons (**Supports Figure 1)**. **c,** Example traces of EEG/EMG and EEG delta power of SD/RS-tagged and control (Off DOX, no SD/RS) *Vgat-PFC-Activity-Tag-ChR2-EYFP* and *Vgat-VC-Activity-Tag-ChR2-EYFP* mice before, during and after light stimulation 10 and 20 Hz over a 50-min period (blue shading). (**Supports Figure 1)**.

**Supplementary Fig. 2.**
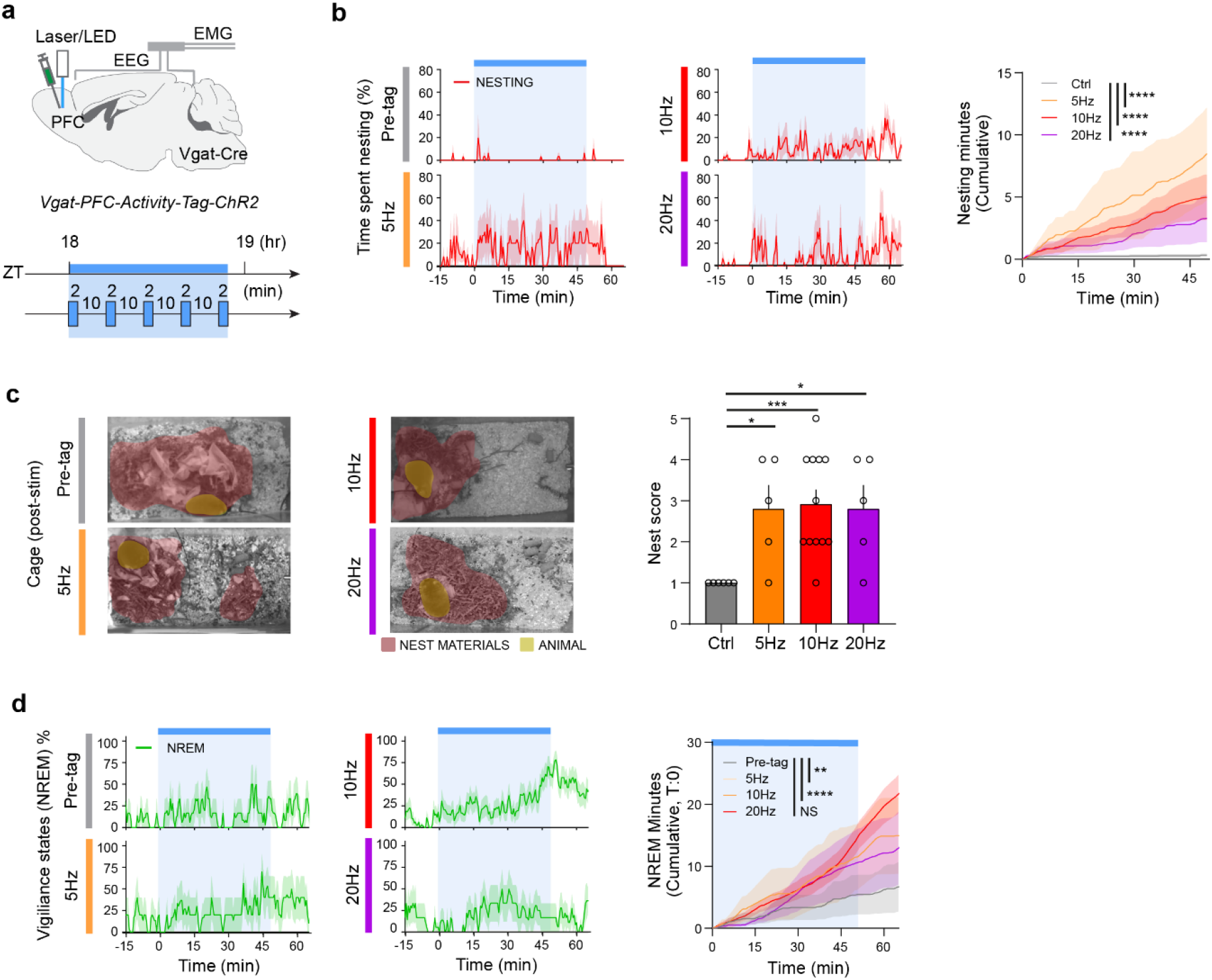
(supports Fig. 1). Following SD/RS-tagging in *Vgat-PFC-Activity-Tag-ChR2-EYFP* mice, increase in nesting and NREM sleep is maximal with 10 Hz opto-stimulation. **a**, Summary of opto-stimulation protocol: repetitive blocks of 2-mins stimulation at either 5, 10 or 20 Hz were given with 10 mins intervals over a 50-min block, starting from ZT 18, which is about 6 hours before lights on. **b**, Time course of nesting activity (time spent nesting, cumulative nesting) of opto-stimulated SD/RS-tagged *Vgat-PFC-Activity-Tag-ChR2-EYFP.* Ctrl (5 mice), PFC (5 mice) each with multiple trials; two-way RM ANOVA, interaction on stimulation group with Ctrl group over time with Bonferroni correction, *F*_99, 792_ = 3.411, p = 8.89E-22 (Ctrl (5 trials) vs 5Hz (5 trials)), *F*_99, 1584_ = 2.465, p = 6.47E-13 (Ctrl (5 trials) vs 10Hz (13 trials)), *F*_99, 891_ = 1.910, p = 1.05E-06 (Ctrl (5 trials) vs 20Hz (6 trials), NS: P>0.05, ****p < 0.0001. Mean (line) ± SEM (shading). Blue shade: Opto-stimulation period. **c,**Pictures of scattered nesting material and nests of *Vgat-PFC-Activity-Tag-ChR2-EYFP* mice after opto-stimulation at either 5, 10 or 20 Hz. Nest material is shaded rose; mouse is shaded olive; and nest scores after opto-stimulation at either 5, 10 or 20 Hz. Ctrl (5 mice), PFC (5 mice) each with multiple trials; Unpaired KS test, D = 0.800, p = 0.0152 (Ctrl (6 trials) vs 5Hz (5 trials)), D = 0.9167, p = 0.004 (Ctrl (6 trials) vs 10Hz (12 trials)),D= 0.800, p = 0.0152 (Ctrl (6 trials) vs 20Hz (5 trials)), NS> 0.05, *p < 0.05, **p < 0.01. Mean ± SEM. **d**, Time course of NREM sleep (cumulative sleep) of opto-stimulated SD/RS-tagged *Vgat-PFC-Activity-Tag-ChR2-EYFP* mice. Ctrl (5 mice), PFC (5 mice) each with multiple trials; two-way RM ANOVA, interaction on stimulation group with Ctrl group over time with Bonferroni correction, *F*_99, 792_ = 3.411, p = 8.89E-22 (Ctrl (5 trials) vs 5Hz (5 trials)), *F*_99, 1584_ = 2.465, p = 6.47E-13 (Ctrl (5 trials) vs 10Hz (13 trials)), *F*_99, 693_ = 0.9631, p = 0.5823 (Ctrl (5 trials) vs 20Hz (6 trials), NS: P>0.05, ****p < 0.0001. Mean (line) ± SEM (shading). Blue shade: Opto-stimulation period.

**Supplementary Fig.3.**
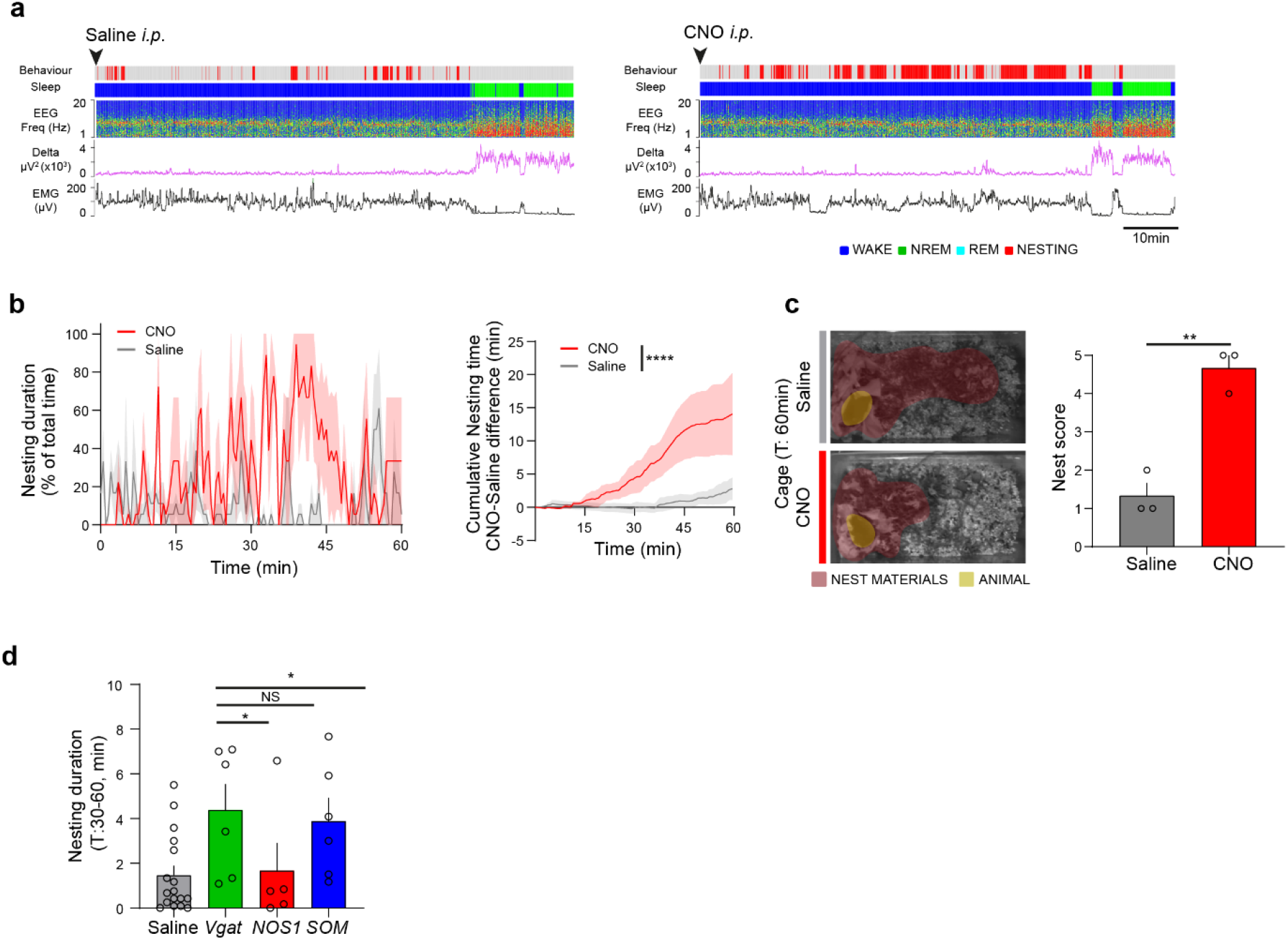
(supports Figs. 1 & 2). **a,** Example traces of EMG and EEG and sleep and behavioral scoring (wake, NREM, REM sleep and nesting) from unilateral chemogenetic SD/RS-tagged *Vgat-PFC-Activity-Tag-hM3Dq-mCherry* mice after Saline and CNO *i.p.* injection. **b,** Time course of nesting activity (nesting duration, cumulative nesting time CNO-saline difference) of SD/RS-tagged *Vgat-PFC-Activity-Tag-hM3Dq-mCherry* mice after Saline and CNO injection. T:0 is *i.p.* injection time point. Saline (3 mice), CNO (3 mice); two-way RM ANOVA, interaction between saline and CNO group over time with Bonferroni correction, *F*_119, 476_ = 4.916, p = 6.852E-36. ****p < 0.0001. Mean (line) ± SEM (shading). **c.** Pictures of scattered nesting material and nests of *Vgat-PFC-Activity-Tag-hM3Dq-mCherry* mice 1 hour after *i.p.* injection. Nest material is shaded rose; mouse is shaded olive; and nest scores. **d.** Time spend Nesting between 30 min and 1 hour (T: 30-60 min) after *i.p.* injection of *Vgat-PFC-Activity-Tag-hM3Dq-mCherry, NOS1-PFC-Activity-Tag-hM3Dq-mCherry and SOM-PFC-Activity-Tag-hM3Dq-mCherry* mice (Saline against CNO 5mg/kg). Saline (a group of Vgat, NOS1, SOM: 17 mice), Vgat (6 mice), NOS1 (5 mice), SOM (6 mice); Unpaired Mann-Whitney test, U = 16, p = 0.012 (saline vs Vgat-CNO), U = 41.5, p = 0.9546 (saline vs NOS1-CNO), U = 18, p = 0.0189 (saline vs SOM-CNO), NS> 0.05, *p < 0.05. Mean ± SEM.

**Supplementary Figure 4.**
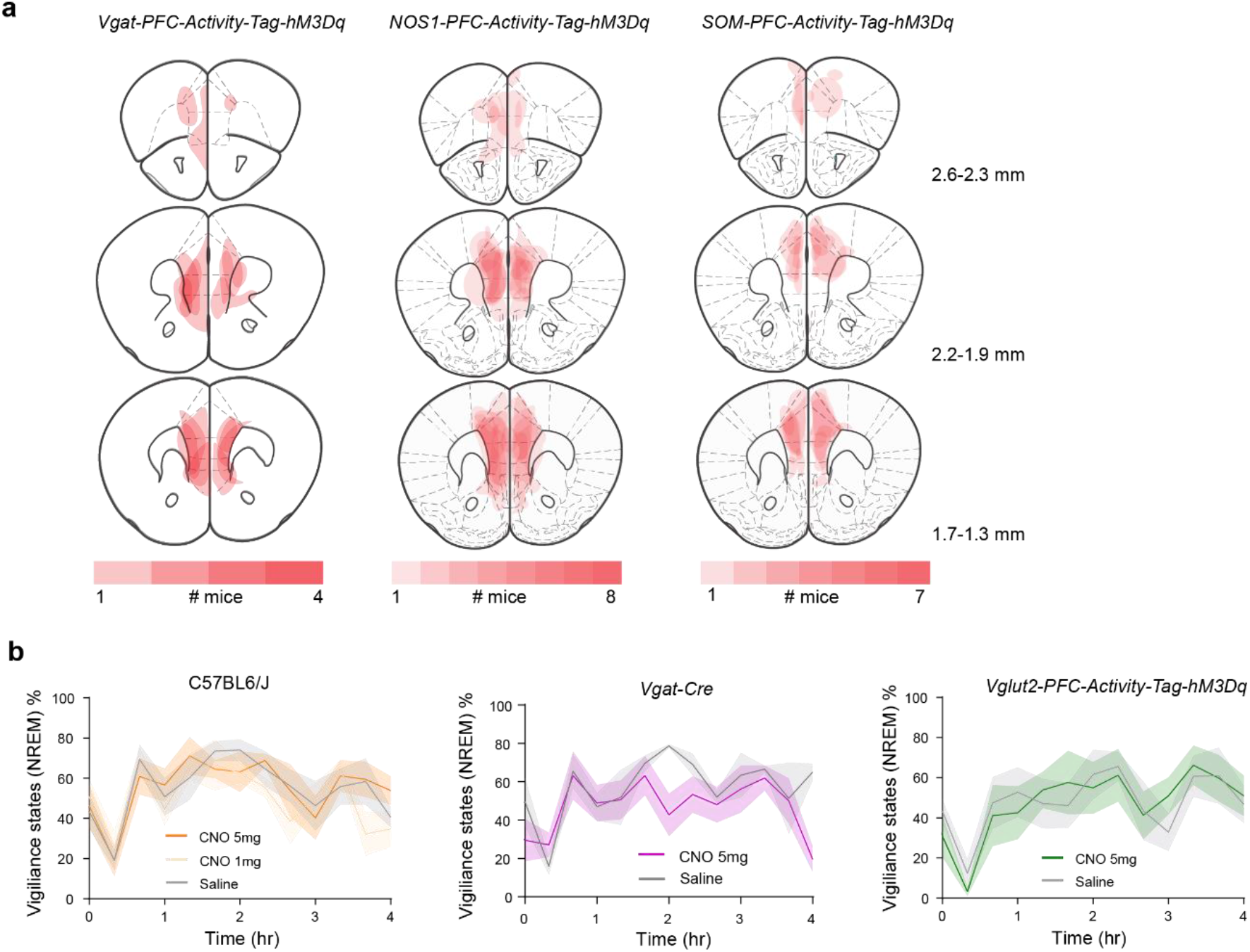
(supports Fig. 2) **a,** Map of hM3Dq-mCherry gene expression induced following SD/RS in *Vgat-PFC-Activity-Tag-hM3Dq-mCherry, NOS1-PFC-Activity-Tag-hM3Dq-mCherry and SOM-PFC-Activity-Tag-hM3Dq-mCherry* mice. Photomicrograph inset shows tagged neurons in the PFC. The schematic diagrams summarize the extent of gene expression the PFC (the intensity of the red indicates the extent of overlap (overlay) of gene expression between animals). **b,** CNO controls. CNO given i.p. at either 5 mg/kg or 1 mg/kg does not induce NREM sleep above baseline in either C57Bl6/J, no AVV injected *Vgat-Cre* and SD/RS-tagged *Vglut2-PFC-Activity-Tag-hM3Dq-mCherry* mice (CNO given at ZT 18). C57Bl6/J (5 mice), *Vgat-Cre* (6 mice), *Vglut2-Cre* (5 mice).

**Supplementary Figure 5.**
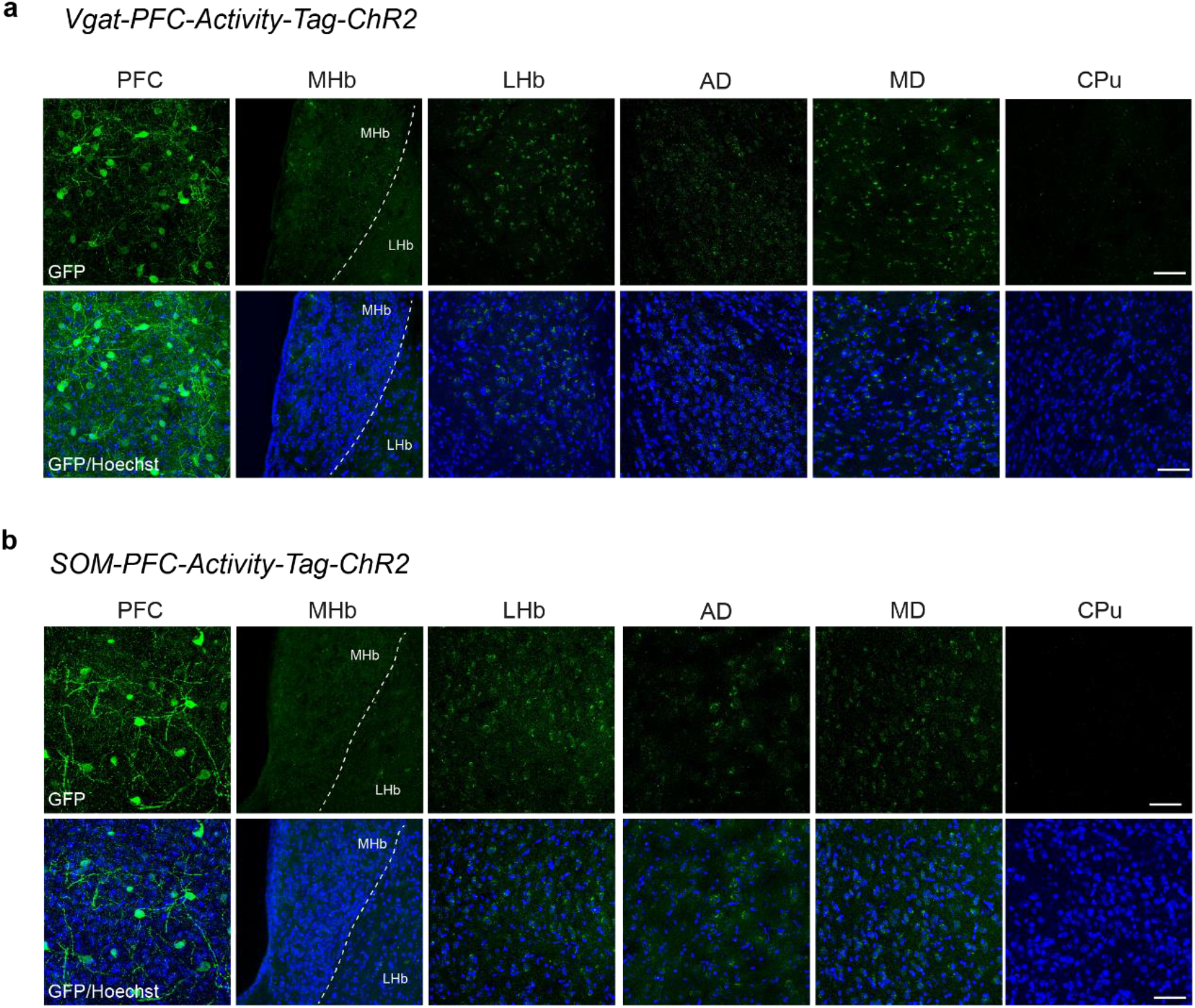
(supports Fig. 4) **a,** *Vgat-PFC-Activity-Tag-ChR2-EYFP* mice following SD/RS tagging: photographs of *ChR2-EYFP-*positive soma and fibers in the PFC, MHb, LHb, AD, MD, and CPu. Scale bar, 50 μm **b,** *SOM-PFC-Activity-Tag-ChR2-EYFP* mice following SD/RS tagging: photographs of *ChR2-EYFP-*positive soma and fibers in the PFC, MHb, LHb, AD, MD, and CPu. Scale bar, 50 μm

